# The microbial community of rust layer biofilm was driven by seawater microbial community

**DOI:** 10.1101/2023.11.03.565569

**Authors:** Shengxun Yao, Junxiang Lai, Congtao Sun, Zihan Pan, Maomi Zhao, Jizhou Duan, Baorong Hou

**Author notes:** Address correspondence to Maomi Zhao,; Jizhou Duan,.

## Abstract

Microbiologically influenced corrosion (MIC) accounts for approximately 20% of the total corrosion-related losses worldwide, causing significant economic damage each year, particularly in Marine environments. However, there are still no truly effective and eco-friendly protection solutions against MIC, among which the incomplete understanding of the microbial biofilm development on metallic surface is a key limitation. Using 16S rRNA and ITS sequencing, we studied bacterial and fungal communities in rust layer biofilm and seawater. The results showed that Proteobacteria, Cyanobacteria and Bacteroidota were the dominant bacterial phyla, and Ascomycota and Basidiomycota were the dominant fungal phyla both in the rust layer biofilm and seawater. Bacteria like Erythrobacter and Aquimarina, as well as fungi like Aspergillus and Acremonium were were notable microorganisms in the rust layer biofilm. Source analysis revealed differences between biofilm and seawater communities, with 23.08% bacterial and 21.48% fungal communities originating from seawater. Stochastic processes governed biofilm and seawater microbial communities, and network analysis showed coexistence and interaction among bacteria and fungi.

**IMPORTANCE:** The composition and source analysis of bacterial and fungal communities in the rust layer and seawater were studied, and the interaction of bacteria and fungi in the rust layer was discussed for the first time. Based on these findings, we provided a framework to explain the observed characteristics of microbial communities in rust layer biofilm and presented key evidence supporting the relationship between different microbial structures and interactions with metal corrosion. These findings, from the perspective of microbial ecology, provide a theoretical foundation for studying microbial corrosion in marine environments.

---

With the development of marine resources around the world, a large number of facilities and equipment have been put into the development of marine economy, such as all kinds of ships, marine exploration facilities, drilling platforms, oil pipelines, offshore wind power facilities and so on. Most of these facilities and equipment are made of carbon steel, stainless steel and other metal materials, which are prone to corrosion in the marine environment (1). It is estimated that the economic losses caused by corrosion in the world every year account for about 3% of the GDP of that year, among which, the losses caused by microbiologically influenced corrosion (MIC) account for about 20% of the total corrosion, and also cause billions of yuan economic losses to the world every year (2), even serious safety accidents. In the process of exploitation and utilization of marine resources, MIC is still an unsolved problem, which need to be paid attention to and solved. There are complex micro fouling biofilms on almost any flooded surface that affect the metal corrosion process through the metabolic activity of microorganisms within the biofilm (3). Therefore, it is a very important topic to analyze the ecological mechanism of microbial community in steel rust biofilm driven by seawater environmental microbial population, and its inherent corrosion mechanisms from the perspective of microbial film.

MIC is the phenomenon of metal corrosion or damage caused by microorganisms adhering to the surface of the metal material, changing the physicochemical conditions of the metal surface through various metabolic reactions and biofilm deposition, etc.(4). Numerous marine microorganisms are capable of causing MIC, including fungi, archaea, algae and different species of marine bacteria (2). As early as the mid-19th century to the 1920s, people began to notice the involvement of microbes and their influence on the corrosion process, and the importance of sulfur reducing bacteria (SRB) in MIC was determined. With the development of interdisciplinary integration and the progress of chemical and electrochemical analysis technology, microbial analysis technology and material characterization technology, the mechanism of MIC has been further explored, and many important microbiological corrosion mechanisms have been put forward (2). Research on MIC primarily focuses on the mechanisms involved in it. MIC is a complex biological and chemical process, where the metabolic activities and byproducts of bacteria and other microorganisms impact the corrosion rate (5). By studying the mechanisms of MIC, researchers have proposed various theories, such as hydrogenase cathodic depolarization theory, metabolic product corrosion theory, concentration cell theory, and extracellular electron transfer theory, particularly in the case of sulfate-reducing bacteria (6). In recent years, concepts like microbially catalyzed corrosion, electroactive microorganisms, and electrochemical active biofilms have emerged. Researchers believe that microbial communities provide energy for cell growth and survival through the transfer of electrons between reducing and oxidizing agents, and this extracellular electron transfer within biofilms play a crucial role in the corrosion process (7).

Bacterial communities within biofilms have received particular attention (8), especially sulfate reducing bacteria (SRB), which is considered associated with most microbial influenced corrosion under anaerobic conditions (9). Researches show that, SRB and its metabolites have cathodic depolarization, which can accelerate hydrogen penetration and assist pitting corrosion, leading to stress corrosion cracking of 980 steel (10); *Desulfovibrio vulgaris* can accelerate the corrosion of duplex stainless steel in eutrophication artificial seawater with a pH 7 to 9 (11); SRB and cathodic protection together can promote the sulfide produced by SRB to penetrate the coating, causing the coating degradation and accelerating the corrosion of the matrix steel (12); and a bridge in Florida, caused severe local corrosion due to degradation of steel pile coating and SRB growth (13). The bacteria associated with MIC under aerobic conditions such as marine *Pseudomonas aeruginosa* also have received particular attention, for it is not only the lead bacterium of biofilm formation, but also has a strong influence in metal MIC. Researches show that, *Pseudomonas aeruginosa* biofilm can significantly reduce the corrosion resistance and accelerate the pitting process of stainless steel (2707 HDSS, 2205 DSS) (14, 15), make high entropy alloy (HEA) passivation film weak and thin (16), and accelerate the corrosion of pure titanium (Ti) metal (17). Some other bacterial communities associated with MIC of metal materials have also received attention and research, such as, iron oxidizing bacteria, iron reducing bacteria and acid-forming bacteria (18, 19).

Fungi is another type of corrosive microorganism that received particular attention. Fungi have a strong ability to thrive on the surface of aluminum alloy in a moist environment. This contributes to the development of more intense corrosion on aluminum alloys (20). In the research progress of the effect of fungi on metal corrosion, the presence of fungi on metal surfaces significantly impacts the strength and durability of the metals. The growth of fungi is influenced by the secretion of metabolites, which allow them to adapt to different environmental and nutritional conditions. While there is limited information available regarding fungi’s ability to adapt to metal surfaces, it is observed that most fungi alter the composition of metals as they grow, engaging them in functional growth and metabolism. This change in composition and color serves as evidence that fungi have penetrated the metal surfaces and utilize them to meet their nutritional requirements, leading to corrosion (21). Numerous studies in recent years have demonstrated the significant influence of fungal communities on the corrosion process. Whether it is fungal-induced atmospheric iron corrosion in indoor environments (22), corrosion of post-tensioned cables (23), or the corrosion of aluminum alloy 7075 induced by marine *Aspergillus terreus* under conditions of absence organic carbon (24), fungi have been found to play a crucial role. The results of these studies highlight the importance of fungi in promoting the corrosion process and underscore the need to acknowledge the impact of fungal communities on corrosion.

Current research on MIC mainly focuses on the corrosion mechanisms of single bacteria and other microbes, which often exist in the form of biofilms in the actual environment. The corrosion mechanism of single species cannot fully explain the corrosion phenomenon in the actual environment. Due to the complexity and variation of marine environment, the microbial community which forming of biofilm is affected by different physicochemical environmental conditions in the areas which it is located, thus affecting the corrosion characteristics of steel materials. Therefore, it is necessary to explore the composition features of microbial communities and their influence on metal corrosion from the perspective of biofilm. Therefore, this project is conducive to understanding the corrosion behavior of metal materials by microbial community in marine environment by studying the composition characteristics of bacteria and fungi in the rust layer biofilm of typical coastal areas in Guangxi and its relationship with bacteria and fungus in seawater environment. Meanwhile, it provides data support for the prevention and control of microbial influenced corrosion. The purpose of this study was to investigate the structural characteristics of microbial communities in the rust layer biofilm and seawater in the survey area, as well as the correlation between them.

## RESULTS

### Environmental factors detection

The physical and chemical properties of seawater samples collected from the nine sampling sites were measured in situ and in lab (Table 1). The seawater samples exhibit a broad spectrum of dissolved oxygen (DO) concentrations, varying from 7.52 to 13.31 mg·L^-1^, with an average value of 10.97 ± 2.10 mg·L^-1^. The seawater pH, temperature and salinity levels exhibited a range of 7.38 to 8.87, 27.70 to 31.23 ℃ and 27.53‰ to 32.71‰, with an average value of 8.31 ± 0.52, 29.56 ± 1.03 ℃ and 30.54‰ ± 1.8‰, respectively. Additionally, the ranges of NH_4_^+^-N, NO_3_^-^-N, NO_2_^-^-N and PO_4_^3-^-P were 0.02-0.23, 0.03-0.29, 0.01-0.04, and 0.02-0.58 mg·L^-1^, and their mean values were 0.12 ± 0.07, 0.13 ± 0.08, 0.02 ± 0.01, 0.26 ± 0.16 mg·L^-1^, respectively.

**Table 1.** The physical and chemical properties of seawater samples.

| Physical and chemical Properties | Statistic analysis |  |  |
| --- | --- | --- | --- |
| | Min | Max | Average ± SD |
| pH | 7.38 | 8.87 | $8.31 \pm 0.52$ |
| Salinity ‰ | 27.5 | 32.7 | $30.5 \pm 1.8$ |
| Temperature | 27.7 | 31.2 | $29.5 \pm 1.0$ |
| DO/ mg·L<sup>-1</sup> | 7.52 | 13.31 | $10.97 \pm 2.1$ |
| NH<sub>4</sub><sup>+</sup>-N/ mg·L<sup>-1</sup> | 0.02 | 0.23 | $0.12 \pm 0.07$ |
| NO<sub>3</sub><sup>-</sup>-N/ mg·L<sup>-1</sup> | 0.03 | 0.29 | $0.13 \pm 0.08$ |
| NO<sub>2</sub><sup>-</sup>-N/ mg·L<sup>-1</sup> | 0.01 | 0.04 | $0.02 \pm 0.01$ |
| PO<sub>4</sub><sup>3-</sup>-P/ mg·L<sup>-1</sup> | 0.02 | 0.58 | $0.26 \pm 0.16$ |

### Microbial comzzposition and diversity in the rust layer biofilm and seawater

A total of 839,954 effective tags of bacteria and 808,814 effective tags of fungi were obtained after filtering the raw data of 18 samples for quality and chimera, and 2,467 OTUs of bacteria and 1,706 OTUs of fungi were identified. All the OTUs were further classified into 38 bacterial phyla and 9 fungal phyla based on the effective tags. To assess and compare the bacterial diversity between the rust layer biofilm and seawater, α-diversity indices such as the Shannon and Chao1 indices were calculated using the OTUs relative abundance of each sample. The bacterial α-diversity showed that the bacterial Shannon index of rust layer biofilm (3.69 ± 0.75) was higher than that of seawater (3.43 ± 0.47) (Fig. 1A), whereas the Chao1 index was lower in rust layer biofilm (267 ± 137) than that of seawater (326 ± 54) (Fig. 1B). For the diversity index of fungi, the Shannon index of fungi in rust layer biofilm (2.04 ± 1.02) was lower than that in seawater (2.79 ± 1.12) (Fig. 1C), and the Chao1 index of rust layer biofilm (261 ± 146) was significantly lower than that of seawater (611 ± 145) (Fig. 1D).

**Fig 1.**
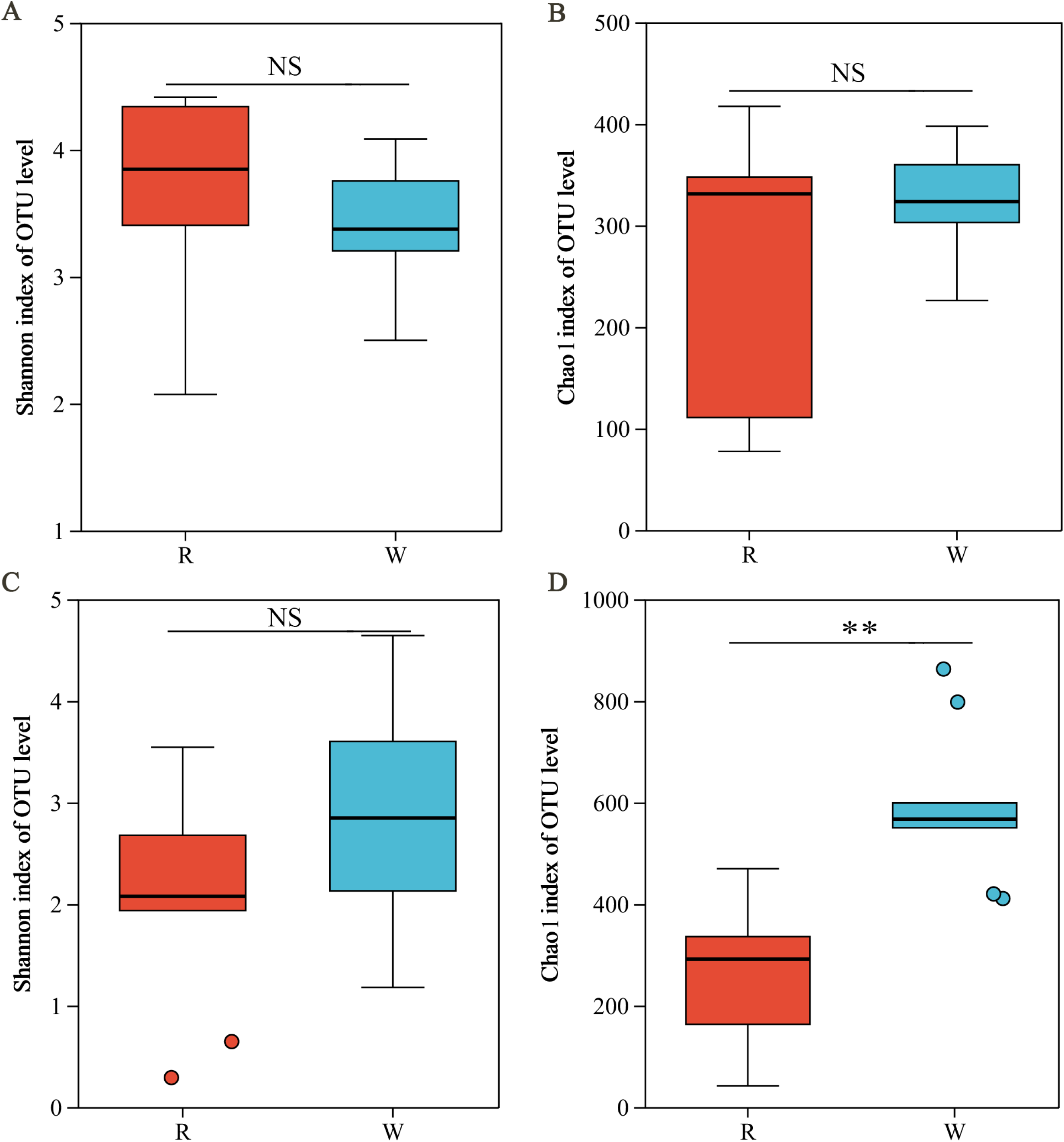
Comparison of microbial diversity. Statistical significance of the α-diversity indices between the rust layer biofilm and seawater were based on the Student’s *t*-test (**: *P* < 0.01). Student’s *t*-test was used to analyze the bacterial Shannon index (A) and the Chao1 index (B), and the fungi Shannon index (C) and the Chao1 index (D) between the rust layer biofilm and seawater (\**P* < 0.05; \*\**P* < 0.01).

In the rust layer biofilm and seawater, Proteobacteria, Cyanobacteria, Bacteroidota, and Actinobacteriota were the dominant bacterial phyla, and Chloroflexi, Firmicutes, and Desulfobacterota were another three dominant bacterial phyla in the rust layer biofilm. Proteobacteria was the most abundant in rust layer biofilm (49.42%), followed by Cyanobacteria (15.10%), Bacteroidota (13.34%) and Actinobacteriota (10.56%), while Cyanobacteria was dominant in seawater (52.56%), followed by Proteobacteria (34.50%) and Bacteroidota (9.06%) (Fig. 2A), indicating that the dominant phyla observed were similar in the two habitats. For fungal communities, unclassified_k Fungi (57.44%), Ascomycota (36.64%) and Basidiomycota (5.85%) was dominant in the rust layer biofilm while unclassified_k Fungi (93.57%) and Ascomycota (5.80%) was more abundant in seawater (Fig. 2B).

**Fig 2.**
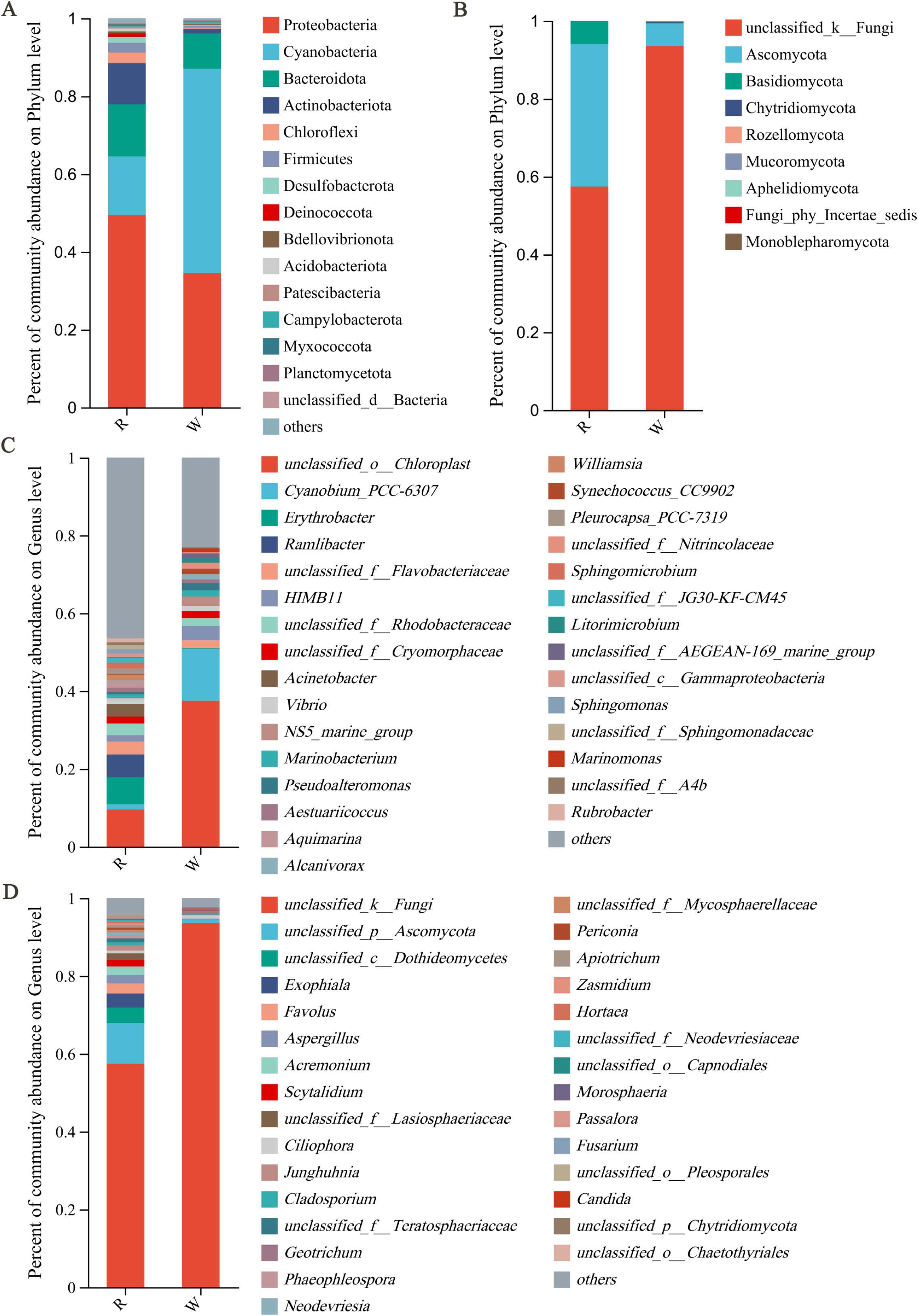
Microbial profiles based on data from sequencing the 16S rRNA and ITS gene amplicon. The top 15 relative abundances of bacterial (A) and fungal (B) phyla in the rust layer biofilm and seawater. The top 30 relative abundances of bacterial (C) and fungal (D) genera in the rust layer biofilm and seawater.

The top 10 bacterial genera in rust layer biofilm include the following: *unclassified_o Chloroplast* (46.46%), *Erythrobacter* (9.53%), *Ramlibacter* (6.98%), *unclassified_f Flavobacteriaceae*(3.34%), *Acinetobacter* (3.20%), *unclassified_f Rhodobacteraceae* (2.99%), *unclassified_f Cryomorphaceae* (1.79%), *HIMB11* (1.67%), *Aquimarina* (1.67%) and *Williamsia* (1.52%). The top 10 bacterial genera in seawater include the following: *unclassified_o Chloroplast* (37.44%), *Cyanobium_PCC-6307* (13.46%), *HIMB11* (3.66%), *NS5_marine_group* (2.44%), *unclassified_f Rhodobacteraceae*(2.04%), *unclassified_f Flavobacteriaceae* (2.02%), *Pseudoalteromonas* (1.86%), *unclassified_f Cryomorphaceae* (1.71%), *Marinobacterium* (1.57%) and *unclassified_f Nitrincolaceae* (1.45%) (Fig. 2C). At the genus level, the major fungal genera in the rust layer biofilm are *unclassified_k Fungi* (57.44%), *unclassified_p Ascomycota* (10.44%), *unclassified_c Dothideomycetes* (3.99%), *Exophiala* (3.62%), *Favolus* (2.60%), *Acremonium* (2.21%), *Aspergillus* (2.16%), *Scytalidium* (1.71%), *unclassified_f Lasiosphaeriaceae* (1.64%) and *Junghuhnia* (1.40%). In seawater, *unclassified_k Fungi* (93.57%) constitutes the vast majority, while *unclassified_p Ascomycota* (1.01%) accounts for a smaller proportion (Fig. 2D).

### Microbial similarities and differences between the rust layer biofilm and seawater

PCoA using Bray-Curtis distances revealed significant differences in the bacterial and fungal composition between the rust layer biofilm (Fig. 3A) and seawater (Fig. 3B) samples. The significant differences community structure between rust layer biofilm and seawater were confirmed by ANOSIM (*P* < 0.001). Venn diagrams were generated to compare the microbial composition between the rust layer biofilm and seawater samples. A total of 541 OTUs of bacterial (Fig. 3C) and 657 OTUs of fungi (Fig. 3D) were shared in rust layer biofilm and seawater, and 1,393 unique OTUs of bacterial, 406 unique OTUs of fungi in the rust layer biofilm, and 533 unique OTUs of bacterial, 643 unique OTUs of fungi in seawater samples, respectively.

**Fig 3.**
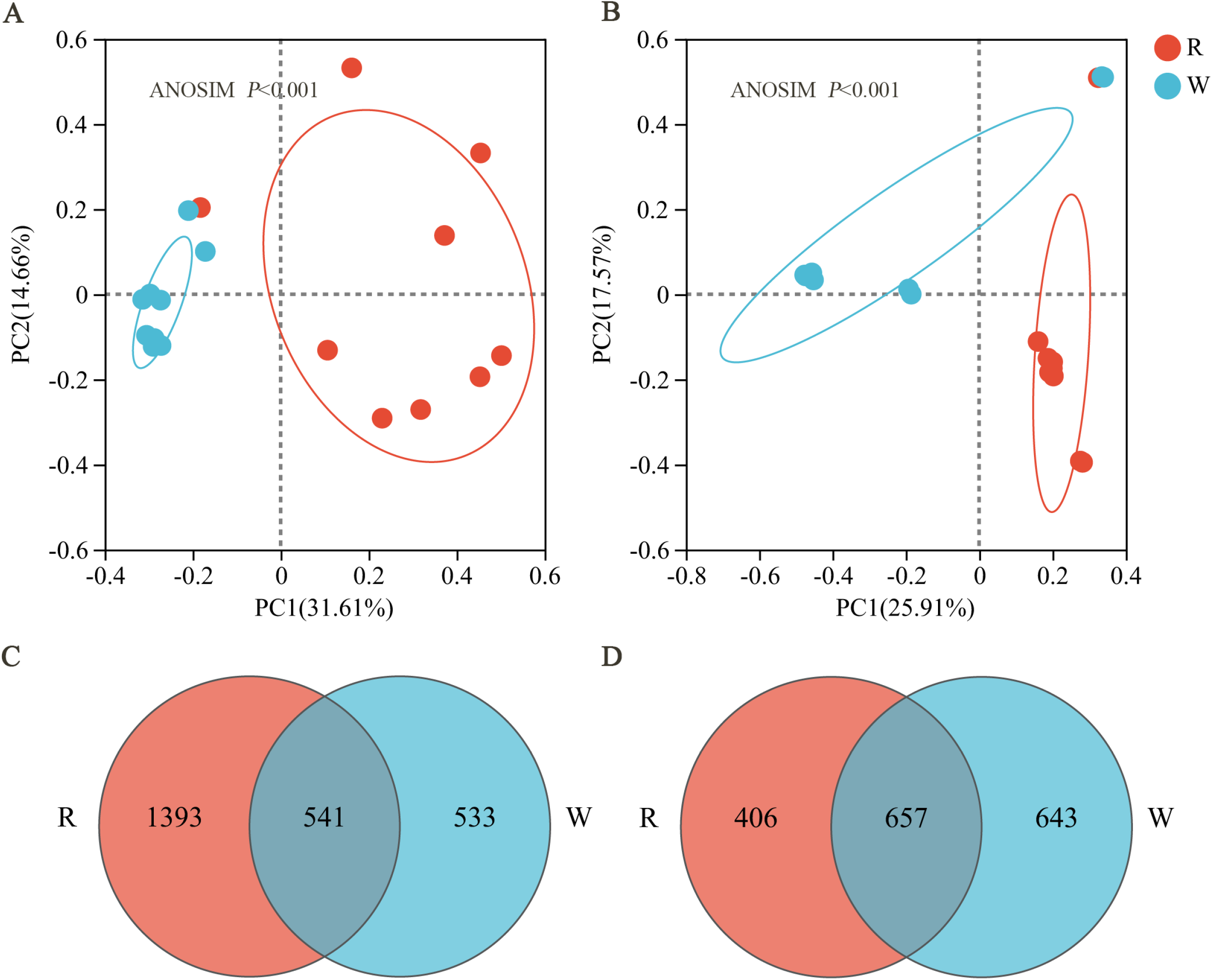
Principal coordinate analysis (PCoA) and ANOSIM analyses of the β-diversity were based on Bray-Curtis distance of the bacterial community structure based on the 16S rRNA gene sequencing (A) and fungal community structure (B) based on the ITS gene sequencing in the rust layer biofilm and seawater. The number of bacterial OTUs (C) and fungal OTUs (D) shared between the rust layer biofilm and the seawater was analyzed by Venn diagram

To determine the genera that exhibited significant differences between the rust layer biofilm and seawater, Student’s t-test was conducted between the two groups. The relative abundances of *Erythrobacter*, *Pleurocapsa_PCC-7319*, *unclassified_f Sphingomonadaceae* and *unclassified_f A4b* bacterial genera were significantly higher in rust layer biofilm than that in seawater, but the relative abundance of the other bacterial genera of the top 15 in seawater were higher in seawater than that in the rust layer biofilm (Fig. 4A). For fungal communities, *Phaeosphaeria* had higher relative abundances in the rust layer biofilm than that in seawater while *unclassified_k Fungi*, *Pseudopithomyces*, *Curvularia*, *unclassified_o GS16*, *Lasiodiplodia*, *Cochliobolus*, *Choanephora*, *Setophaeosphaeria* and *Fusicolla* had lower relative abundances in seawater than in rust layer biofilm (Fig. 4B).

**Fig 4.**
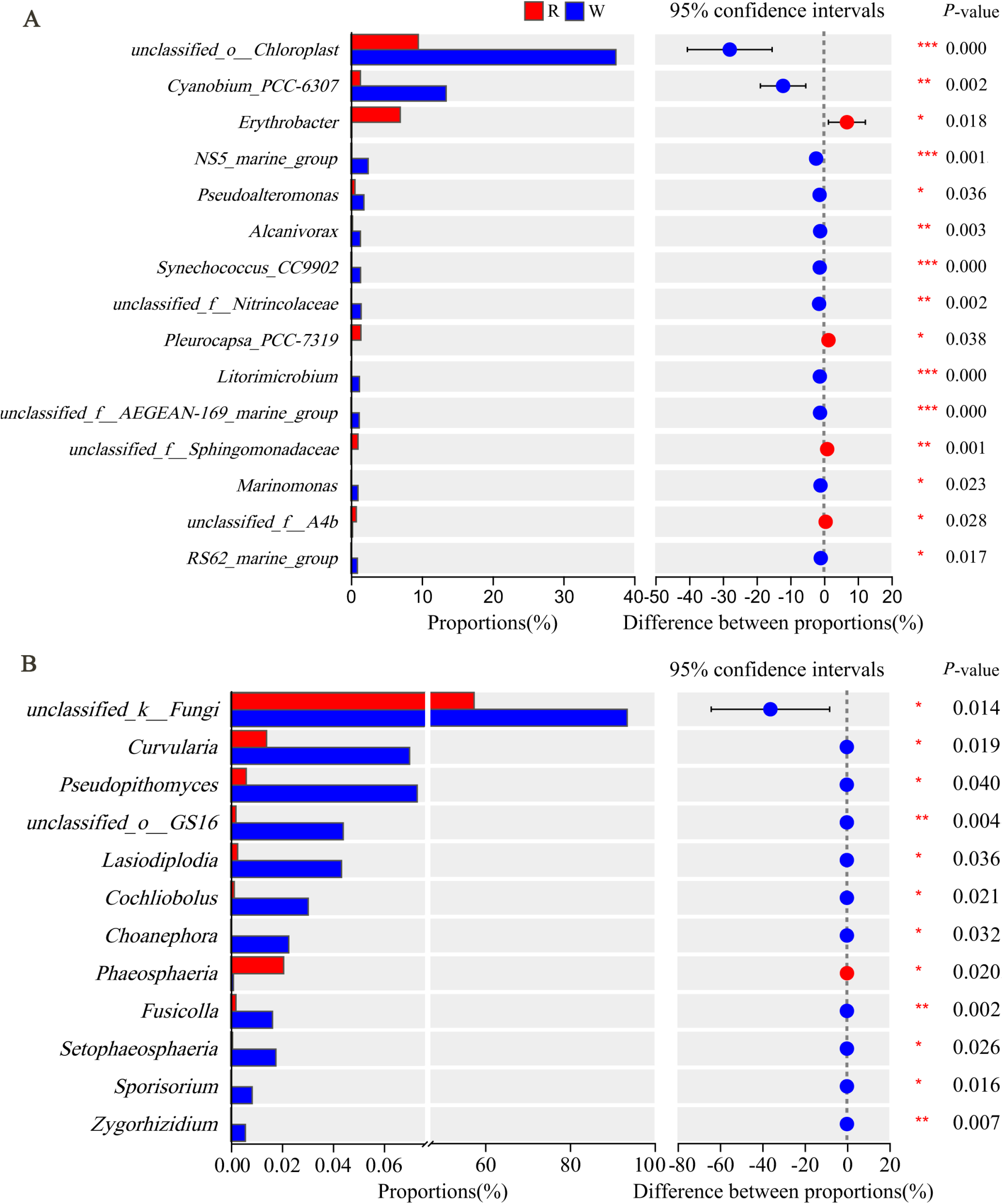
Student’s t-tests revealed distinct bacterial (A) and fungal (B) communities in the rust layer biofilm and seawater at the genus levels (\**P*<0.05, ** *P* < 0.01, *** *P* < 0.001).

### Bacteria and fungi coexist in both rust layer biofilm and seawater

The bacterial networks results suggested that the rust layer biofilm network had 49 nodes connected by 157 edges (Fig. 5A) and the seawater network had 50 nodes connected by 361 edges (Fig. 5B). Based on the network analysis results, it can be observed that the bacterial communities in both the rust layer biofilm and seawater showed relatively high connectivity. The average degree indices of the rust layer biofilm and seawater bacterial communities were 6.41 and 14.44, respectively.This indicates that each bacterial species interacted with several other species within their respective groups. Additionally, the average clustering coefficient values for the rust layer biofilm and seawater was found to be 0.58 and 0.69, respectively. These coefficients suggest that there is a moderate level of clustering or grouping of bacteria within each community. This implies the presence of closely connected subsets of bacterial species within the larger network. Furthermore, the co-associations between bacterial species were predominantly positive in both rust layer biofilm and seawater. With relative positive co-occurrence rates of 70.06% and 65.93% in the rust layer biofilm and seawater, respectively, this suggests a high level of interspecies cooperation or mutualism among the bacterial community members. Overall, the network analysis results indicate that the bacterial communities in both the rust layer biofilm and seawater exhibit complex and interconnected relationships. The high connectivity and positive co-associations suggest the presence of strong interspecies interactions and potentially higher levels of interspecies competitive activities within these communities. Furthermore, the genus of *unclassified_f Alteromonadaceae* (degree = 14), *Sphingomonas* (degree = 12) and *Erythrobacter* (degree = 12) all had high nodes were identified as keystone species in the rust layer biofilm, and *NS5_marine_group* (degree = 28), *Rheinheimera* (degree = 26) were identified as key species in seawater, suggesting their potential roles in maintaining the structure and functionality of the ecological community.

**Fig 5.**
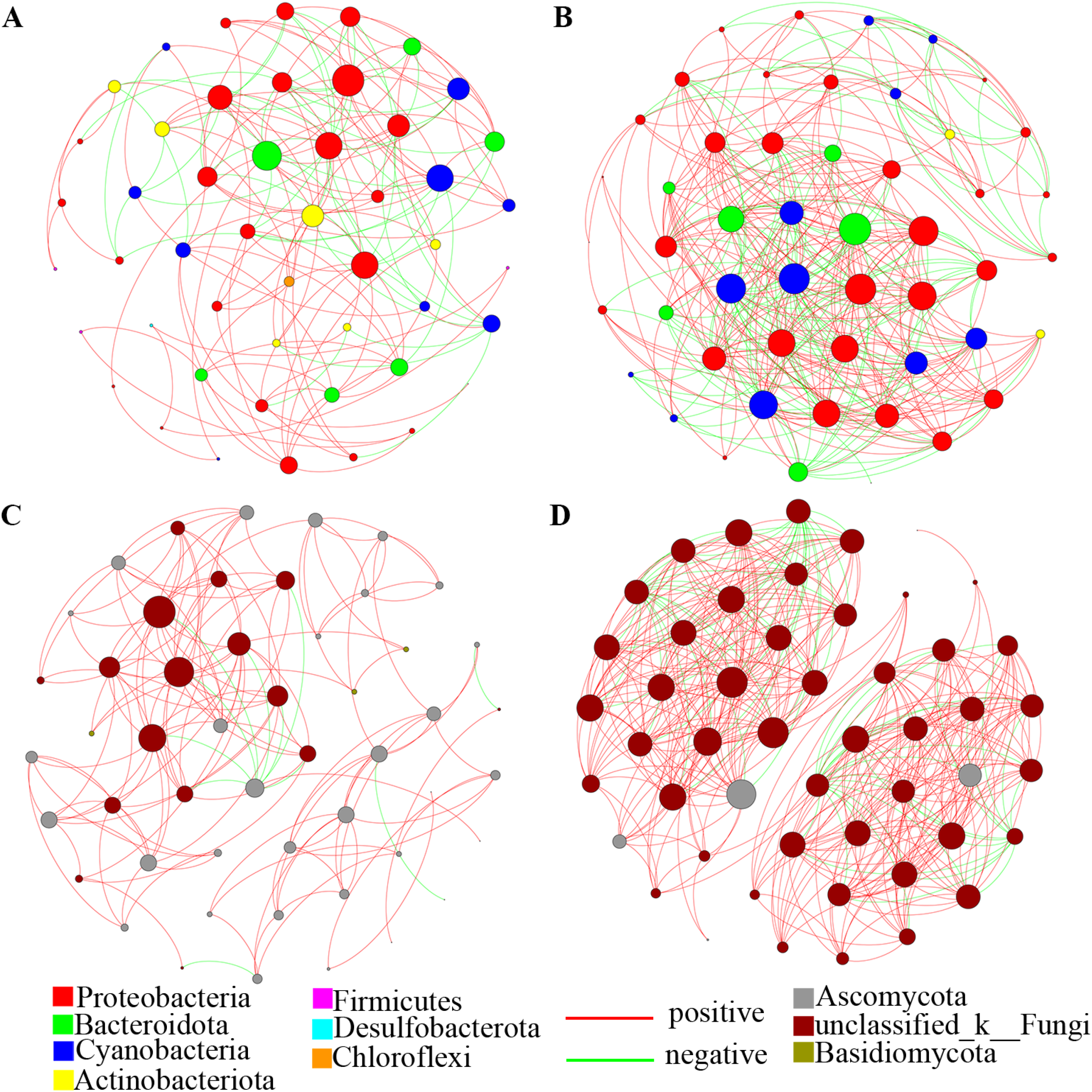
Interspecies interactions between the rust layer biofilm (A) and seawater bacteria (B), and the rust layer biofilm (C) and seawater fungi (D). OTU is represented by each node. The colors of the nodes indicate which OTU is associated with which major phylum. Positive interaction between two individual nodes is indicated by a red edge, while negative interaction is indicated by a green edge.

For fungal networks, the rust layer biofilm network had 50 nodes connected by 146 edges (Fig. 5C) and the seawater network had 50 nodes connected by 417 edges (Fig. 5D). The average degree indices of fungal communities in the rust layer biofilm and seawater were 5.84 and 16.68, and the average clustering coefficient index values were 0.64 and 0.87 and the average path distance were 4.82 and 1.34, respectively. Though some fungi genera had high nodes, include *unclassified_p Ascomycota* (degree = 23) as key species in the rust layer biofilm and in seawater, all were unclassified at genus level.

### Interaction of bacteria and fungi in rust layer biofilm and seawater

Bacteria and fungi coexist and interact with each other in rust layer biofilm (Fig. 6A) and seawater (Fig. 6B). The analysis of the network indicates that the rust layer biofilm network is comprised of 859 nodes, connected by 2,432 edges, and the seawater network consists of 999 nodes, interconnected by a total of 9,848 edges. Moreover, the average degree indices were 5.67 and 19.72 in the rust layer biofilm and seawater, respectively. Notably, a predominantly positive co-occurrence pattern was observed, particularly within the rust layer biofilm. The relative proportion of positive co-expression in the rust layer biofilm and seawater measured at 90.67% and 54.26%, respectively. This suggests a significant coexistence of bacteria and fungi in the rust layer biofilm, indicating an elevated level of interspecific competition activity.

**Fig 6.**
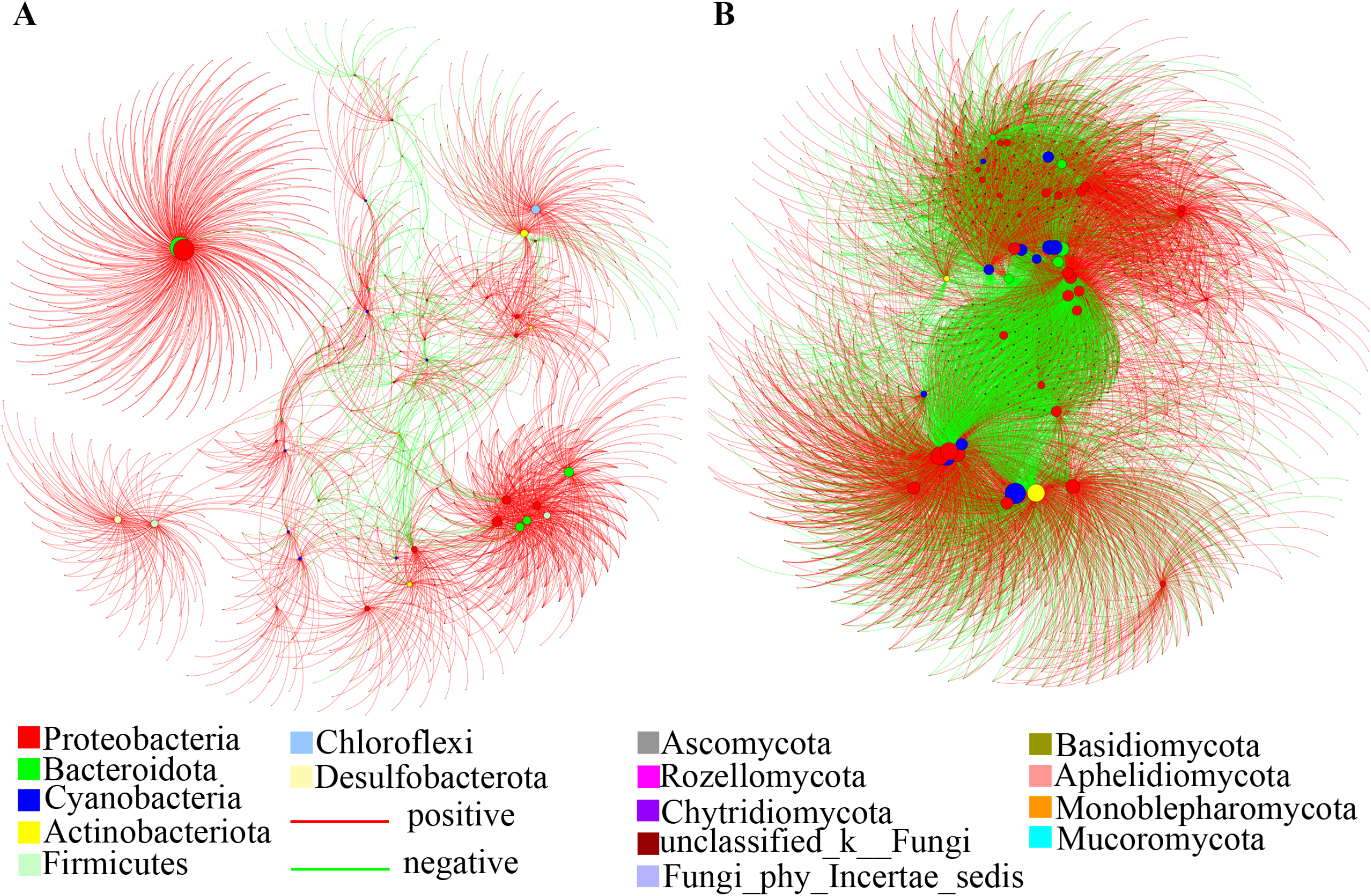
Species-species association networks of bacterial (A) and fungal (B) communities in the rust layer biofilm and seawater.

For bacteria-fungi co-occurrence in the rust layer biofilm, the genus of *Rhodococcus* (degree = 242), *unclassified_f Rhodobacteraceae* (degree = 100), *unclassified_f Flavobacteriaceae* (degree = 109), *Erythrobacter* (degree = 98), *Desulfobacter* (degree = 83) were all keystone species that they have many neighbors in the rust layer biofilm network, which belong to the phyla of Actinobacteriota, Proteobacteria, Bacteroidota and Desulfobacterota. In seawater, the genera of *unclassified_o Chloroplast* (degree = 430), *Synechococcus_CC9902* (degree = 390), *Ascidiaceihabitans* (degree = 378) and *Candidatus_Actinomarina* (degree = 367) were keystone species in the seawater biofilm network, which belong to the phyla of Cyanobacteria, Proteobacteria and Actinobacteriota.

### SourceTracker analysis of contributions of rust layer biofilm and seawater source communities to each other’s communities

In order to test whether the microbial community in seawater plays a decisive role in the microbial community on rust layer biofilm, we used SourceTracker to evaluate the contribution of seawater-originated microbial communities to rust layer biofilm. The results show that the contribution proportion of seawater-originated bacterial communities to the rust layer biofilm was found to be 23.08% (Fig 7A). In contrast, the contribution of the rust layer biofilm to the seawater communities was 76.33% (Fig 7B). The contribution of seawater fungal community to rust layer biofilm was 21.48% (Fig 7C), and the contribution of rust layer biofilm to seawater was 22.26% (Fig 7D). These results suggest that the microbial community of rust layer biofilm was driven by seawater microbial community.

**Fig 7.**
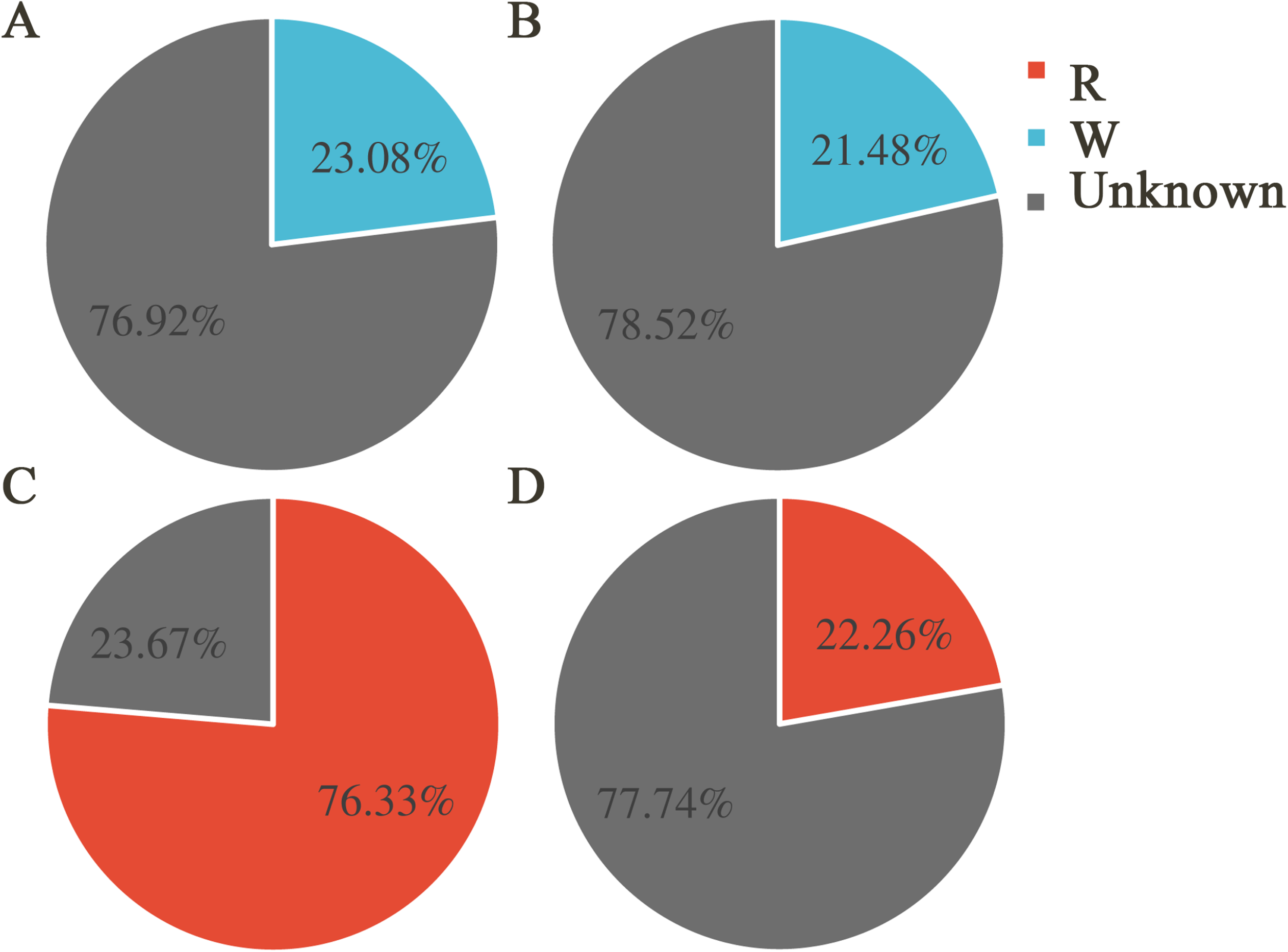
The contribution of seawater bacterial community to rust layer bacterial community (A) and rust layer bacterial community to seawater bacterial community (B) were analyzed by using SourceTracker, as well as the seawater fungi community to rust layer fungi community (C) and rust layer fungi community to seawater fungi community (D).

### The contribution of different ecological processes to the assembly of microbial communities in the rust layer biofilm and seawater

The relative contributions of different ecological processes in the assembly of microbial communities in rust layer biofilm and seawater were assessed using iCAMP. The results revealed that, for bacteria, approximately one-third of the observed changes in the rust layer biofilm community could be attributed to heterogeneous selection (4.74%), homogeneous selection (22.81%), dispersal limitation (30.38%), homogenizing dispersal (2.33%), and drift (39.74%) processes (Fig. 8A). In the seawater, heterogeneous selection accounted for 0.35%, homogeneous selection for 32.21%, dispersal limitation for 10.53%, homogenizing dispersal for 4.39%, and drift for 52.52%. Comparatively, in both the rust layer biofilm and seawater, heterogeneous and homogeneous selection processes made a partial contribution to the bacterial community in the rust layer biofilm (Fig. 8B). Regarding the assembly processes of fungal communities, in the rust layer biofilm, heterogeneous selection accounted for 16.02%, homogeneous selection for 13.45%, dispersal limitation for 38.79%, homogenizing dispersal for 1.35%, and drift for 30.38% (Fig. 8C); in the seawater, heterogeneous selection accounted for 0.84%, homogeneous selection for 42.89%, dispersal limitation for 30.24%, homogenizing dispersal for 2.29%, and drift for 30.24% (Fig. 8D).

**Fig 8.**
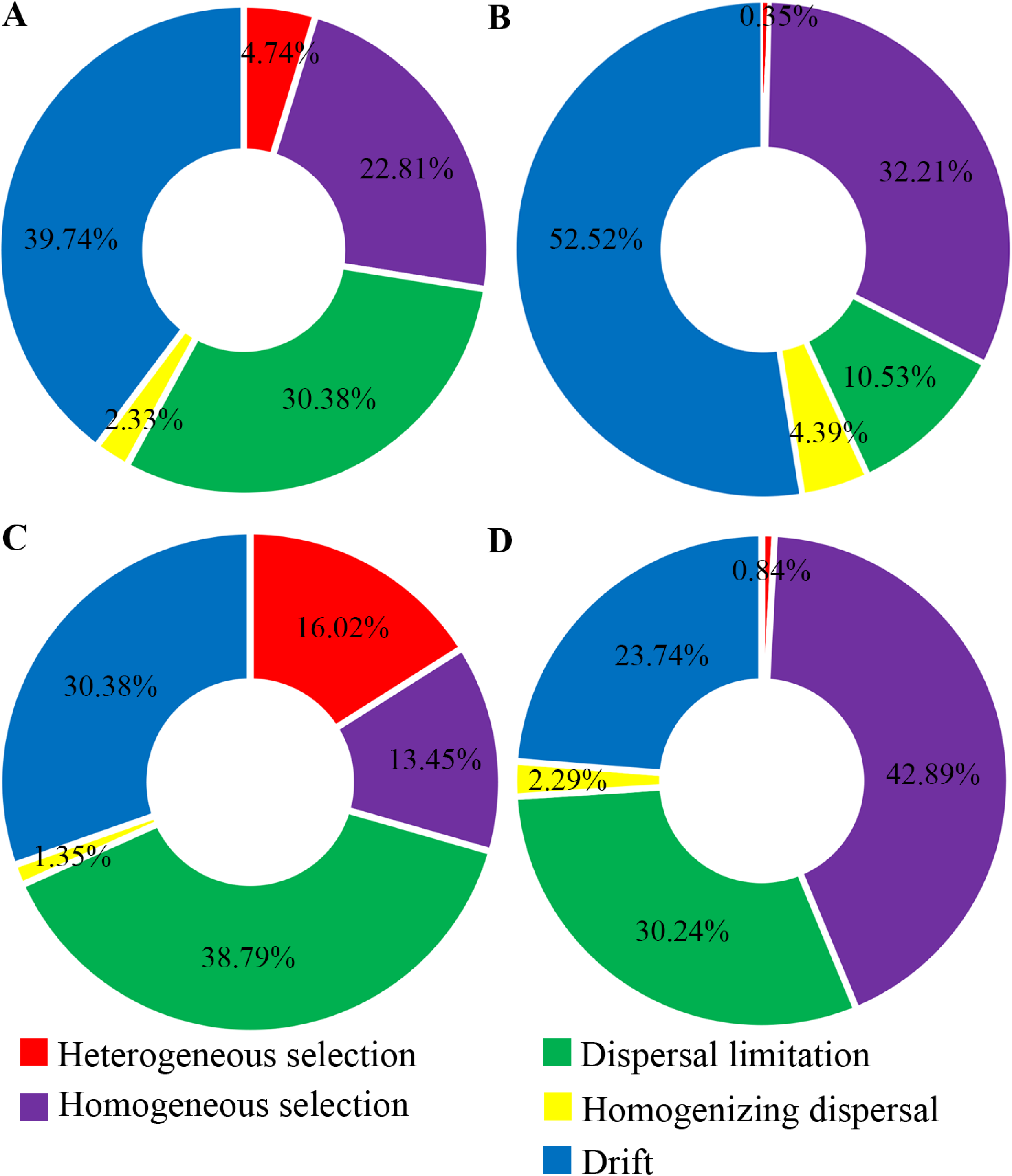
Ecological process analysis of seawater bacteria (A) and rust layer biofilm bacteria (B), as well as ecological process analysis of seawater fungi (C) and rust layer biofilm fungi (D), were conducted.

## DISCUSSION

### Corrosion-related bacteria are enriched in the rust layer biofilm

Many studies on the corrosion of metal alloys in marine environments tend to concentrate on specific species or groups within controlled laboratory conditions, disregarding the involvement of diverse microorganisms in the process. Therefore, there is a lack of understanding regarding the bacterial and fungal communities present and their interactions within rust layer in the natural marine environment. In this research, it was discovered that the rust layer biofilm harbored a diverse range of microbial species, with both bacteria and fungi coexist. The abundance of these microorganisms indicates the potential synergistic or competitive interactions among them during the process of MIC.

A small number of studies have focused on rust layer biofilm microorganisms in the type of metals and found that they can vary widely at the OTU or genus level depending on the location and environmental conditions but tend to be dominated by *Acidithiobacillus ferrooxidans, Leptospirillum ferriphilum, Sulfobacillus thermosulfidooxidans,* and *Acidithiobacillus thiooxidans*, which are known to play a role in the corrosion process by oxidizing metal ions and producing acid, which can further accelerate the corrosion of metals (25). These studies have revealed common conclusions about the bacterial composition of the rust layer biofilm, but the core taxa of the rust layer biofilm are still unknown. In our study, we discovered that although bacterial communities in the rust layer biofilm were highly diverse, in terms of bacterial composition, Proteobacteria, Cyanobacteria, Bacteroidota, Actinobacteriota Chloroflexi, Firmicutes and Desulfobacterota are the major bacterial phyla in the rust layer biofilm, which is consistent with previous studies (26). At the genus level of bacteria, the phototrophic heterotrophic bacterium *Erythrobacter* is a dominant genus in the rust layer biofilm. *Erythrobacter* have been documented that these organisms inhabit tropical marine environments where they have a tendency to attach themselves to marine structures. They have been associated with corrosion attacks on steel alloys and stainless alloys exposed in seawater, such as the Red Sea, Mediterranean Sea, and China South Sea, and have been identified in various marine environments, including ship, drilling platforms and marine structures (27, 28). In a study on the impact of deep-sea bacteria on Cu-Ni-Si alloys, it was discovered that chloride ions disrupted the thin film formed on the alloy surface, thereby accelerating the corrosion rate (29). A separate investigation revealed that deep-sea bacteria are capable of forming a dense biofilm, acting as a protective shield against ongoing corrosion processes (30). These microorganisms possess genes associated with aerobic phototrophy and contain bacterial chlorophyll, enabling them to perform photosynthesis without producing oxygen. They can fix carbon dioxide and participate in heterotrophic processes (31). Moreover, *unclassified_f Rhodobacteraceae* is also abundant in rust layer biofilm. Research has shown that Rhodobacteraceae species are commonly found in the in-situ succession of soft steel communities (32), which are also one of the most diverse and abundant bacterial categories in studies on carbon steel bio-corrosion in seawater (33). They have the ability to colonize newly formed surfaces, and the extracellular polymeric substances (EPS) they produce promote the settlement of other microorganisms, playing an important role in the formation of biofilm (34). Prior research has provided evidence that *Ramlibacter* contains multiple copies of genes regulating nitrogen degradation, nitrogen fixation, and assimilatory nitrate reduction pathways. It dominates several nitrogen metabolism divergent pathways in the microplastic sphere (35), suggesting that the metabolism of such bacteria may be related to the formation of rust layer biofilm. Our research shows that *Aquimarina* had a higher relative abundances in the rust layer biofilm samples, which enhanced corrosion and α-FeOOH formation (36).

### Corrosion-related fungi are enriched in the rust layer biofilm

In recent years, a multitude of studies have unraveled the significant impact exerted by fungal communities on the corrosion process. Whether it involves fungal-induced atmospheric iron corrosion within enclosed spaces (22), the decay of post-tensioned cables (23), or the corrosion of aluminum alloy 7075 triggered by marine *Aspergillus terreus* in conditions of sustained organic carbon deprivation (24), fungi have emerged as pivotal actors. The findings derived from these investigations underscore the indispensable role played by fungi in fostering corrosion, urging us to acknowledge and address the influence wielded by fungal communities on this process. At the genus level of fungi, *Aspergillus* is one of the main fungi in the rust layer biofilm. Multiple research investigations have presented compelling evidence supporting the notion that *Aspergillus* sp., a type of fungus, possesses the ability to accelerate the deterioration process of titanium, carbon steel, and magnesium alloy when exposed to eutrophic environments (37). The production of organic acids by *Aspergillus* sp. has been extensively documented as having detrimental effects on various materials, leading to corrosion and destruction (38). The resilience of *Aspergillus terreus* in the face of organic carbon deficiency enables its ability to establish a biofilm on the surface of Aluminum alloys, thereby hastening the corrosion of aluminum alloys (24). The increased corrosion rate was observed to be a direct result of the accelerated anode and cathode reactions caused by the *Aspergillus niger* biofilm. Moreover, it has been conclusively determined that the corrosion of aluminum alloy is primarily caused by organic acid corrosion induced by the presence of *Aspergillus niger* (39). The corrosion rate of aluminum was significantly affected by the growth and adherence of *Acremonium* sp. (40).

### Bacteria and fungi coexist and interact in the rust layer biofilm

Our research indicated that there is coexistence and interaction between bacteria and fungi in both the rust layer biofilm and the surrounding seawater. Bacteria are identified as the main core species in the coexistence network of bacteria and fungi. Additionally, the key species differ between the rust layer biofilm and seawater. Our study on bacteria-fungi co-occurrence result reveals that the phyla of Actinobacteriota, Proteobacteria, Bacteroidota and Desulfobacterota are dominant in rust layer biofilm, and Cyanobacteria, Proteobacteria and Actinobacteriota are dominant in seawater, which is similar to the findings in other marine environments (41). Exploring the connectivity maintained by modules using intermediacy centrality has been employed in some studies to identify key species in ecosystems (42). Our study shows that bacteria and fungi coexist and engage in interactions within both the rust layer biofilm and seawater which implies that inter-kingdom interactions among microorganisms are significant importance in shaping the diversity, variability, and stability of microbial communities (43). These interactions range from mutualistic to parasitic and involve both contact-dependent and contact-independent mechanisms (44). By analyzing amplicon sequencing data of bacterial and fungal communities, network analysis has revealed strong correlations between the abundance of bacterial and fungal taxa in the rust layer biofilm and seawater. Our research findings indicate that symbiotic interactions between bacteria and fungi were observed in both the rust layer biofilm and seawater. In general, competitive interactions were more widespread (45), as the production of antimicrobials or resource utilization is considered the primary force driving the balance between fungi and bacteria in host niches (46).

### The rust layer biofilm microbe derived from seawater microbe

Our research findings indicated that the contribution proportion of seawater-originated bacterial communities to the rust layer biofilm was found to be 23.08%. In contrast, the contribution of the rust layer biofilm to the seawater communities was 76.33%. One possible explanation for these observations is that, in order to form a rust layer biofilm on the steel surface, the rust layer biofilm selectively recruits bacteria from the aquatic environment with specific functional traits to perform various functions. This is similar to known dominant bacteria in rust layer biofilms, such as sulfate-reducing bacteria and *Pseudomonas aeruginosa* (9, 14, 15). The importance of heterogeneous and homogeneous selection processes increases in the rust layer biofilm bacterial community, promoting the formation of the biofilm. The colonization of bacteria from the aquatic environment into the rust layer biofilm is the result of deterministic processes, but stochastic processes also facilitate the establishment and success of external microbial populations (47). Our source tracking analysis results also support this notion, as they indicate that the bacterial community in the rust layer biofilm is randomly dispersed into the seawater, resulting in a higher contribution of the rust layer biofilm to the seawater microbial community compared to the contribution of the seawater bacterial community to the rust layer biofilm. For fungi, the contribution of seawater fungal community to rust layer biofilm was 21.48%, and the contribution of rust layer biofilm to seawater was 22.26%. Furthermore, the contribution of the seawater fungal community to the rust layer biofilm is comparable to the contribution of the rust layer biofilm to the seawater. This suggests that the establishment of the fungal community in the rust layer biofilm is likely the result of a combination of stochastic and deterministic processes. In the seawater fungal community, homogeneous selection plays a significant role. Homogeneous selection can be seen as the combined effects of host filtering and environmental filtering, ultimately leading to similar community composition characteristics among different individuals or in similar environments (48). Regardless of whether it is the rust layer biofilm or seawater microorganisms, stochastic processes dominate the assembly of bacterial and fungal communities.

The microorganisms present in the rust layer and the seawater were significantly different, but there were still a significant number of commensal microbes between them, suggesting a potential microbial exchange pathway between the two groups. When the microbial communities in seawater enter the steel surface, they change the surface environment through various physiological and biochemical reactions, eventually leading to the formation of rust layer (49). The bacteria in the rust layer biofilm can be divided into two categories, one is naturally existing bacteria from seawater, and the other is bacteria with in situ growth caused by bacteria in seawater (50). The corrosion of bacterial community in seawater in phosphoric acid medium is an important control factor, which can promote the formation of rust layer bacterial community by producing water-soluble products and thus affect the corrosion behavior of steel material (51). Therefore, the seawater microbial community is a key factor in shaping the formation and development of the rust layer biofilm microbial community, and understanding their impact on steel corrosion in the marine environment is of utmost importance.

### Conclusion

In summary, our work has established the relationship between bacterial and fungal interactions in the rust layer biofilm and microbiologically influenced corrosion (MIC). Specifically, significant differences exist between the microbial communities of the rust layer biofilm and seawater. Overgrowth of *Erythrobacter*, *norank_f Rhodothermaceae*, *Aquimarina* bacteria, as well as *Aspergillus* and *Acremonium* fungi on the surface of steel may lead to the occurrence of MIC. The higher relative abundance of these bacteria and fungi within the rust layer biofilm, potentially exacerbating the corrosive process. Furthermore, seawater microorganisms penetrate and colonize the biofilm within the rust layer biofilm, and the interaction between foreign deposited microorganisms and those within the biofilm influences the construction of microbial communities within the rust layer biofilm. The potential reason was that stochastic processes dominate the assembly of the rust layer biofilm and seawater microbial communities. Based on these findings, we provide a framework to explain the observed characteristics of microbial communities in the rust layer biofilm and offer key evidence indicating the relevance of different ecological structures and interactions within these communities to metal corrosion. The results of this study contribute to our understanding of the composition characteristics of microbial communities in both the rust layer biofilm and seawater, as well as the relationship between microbial communities within the rust layer biofilm and the seawater. These research findings provide a theoretical foundation from the perspective of microbial ecology for studying MIC in marine environments

## MATERIALS AND METHODS

### Sample collection and seawater physical and chemical properties measuring

A total of 18 samples (9 rust layer biofilm samples and 9 seawater samples) were collected from Fangcheng Port (108.34° E, 21.62° N), Qinzhou Port (108.61° E, 21.75° N) and Beihai Port (109.10° E, 21.48° N) in November 2022. The rust layer biofilm samples were collected and transferred into a sterile centrifuge tube. Each sample was a mixture of three independent original samples collected from the rust layer at an approximately 5.5 cm^2^ area (26). In order to compare with the microbial community composition in the rust layer, microbes in the surrounding seawater were collected simultaneously to represent microbial in the vicinity seawater, and it was collected by filtering seawater through a 0.22 μm sterile filter membrane. All the samples collected were initially stored in an ice box before transferred to the laboratory, and then kept at -80 ℃ for further analysis. Additionally, some physical and chemical properties of the seawater were measured *in-situ*, including temperature and dissolved oxygen (DO) concentrations with a portable dissolved oxygen meter (JPBJ-608, INESA, China), pH value with a portable pH meter (PHBJ-261L, INESA, China), and salinity levels with a salinometer (DLX-ARH100, DEILIXI, China), while the other properties (NH_4_^+^-N, NO_2_^-^-N, NO_3_^-^-N and PO_4_^3-^-P) were measured in the laboratory with an automatic nutrient analyzer (San++, Skalar, Dutch).

### DNA extraction and 16S rRNA and ITS sequencing

The rust layer biofilm and seawater samples were subjected to DNA isolation using the Bacterial DNA Isolation Kit (Mobio, Carlsbad, CA, USA) and Water DNA Isolation Kit (Omega Bio-tek, Doraville, GA, USA), respectively. The concentration and quality of the DNA samples were determined with a NanoVuePlus spectrophotometer (Thermo Fisher, Massachusetts, USA), and were visualised on a 1% agarose gel. The V4 hypervariable region of the 16S rRNA gene was amplified using the primer pair 338F (5’-ACTCCTACGGGAGGCAGCAG -3’) and 806R (5’-GGACTACHVGGGTWTCTAAT-3’), with random 6-base oligo barcode labelling (52), and the ITS2 gene using the primer pairs ITS3F (5’-GCATCGATGAA GAACGCAGC-3’) and ITS4R (5’-TCCTCCGCTTATTGATATGC-3’) for fungi (53). We generated sequencing libraries using the TruSeq DNA PCR-Free Sample Preparation Kit (New England Biolabs, Massachusetts, USA), which were sequenced using the Illumina MiSeq PE300 platform (Illumina, California, USA).

### Data analysis

The raw data was obtained from the MiSeq PE300 platform, and it was processed as paired-end reads. Each sequence was subsequently assigned to a corresponding sample based on its unique barcode. To eliminate the barcode and primer sequences, the Quantitative Insights into Microbial Ecology (QIIME) software was utilized (54). The short reads were rapidly adjusted to obtain raw tags by pairing and merging paired-end reads (55). To obtain high-quality and clean tags, the QIIME software was employed to filter out low-quality raw tags. The UCHIME algorithm was then utilized to identify and eliminate any chimeric tags, resulting in the successful acquisition of effective tags (56). Uparse was utilized for effective tag analysis leading to the final step of obtaining OTUs. Sequences with a similarity of over 97% were then clustered into the same OUT, and subsequently, annotated according to the OTU (56). Additionally, Classification information for each representative sequence was annotated against the Silva (Release138, http://www.arb-silva.de) bacterial database and the Unite (Release 8.0) fungal database using the Ribosomal Database Project (RDP) classifier with an 80% confidence threshold (57). The abundance of OTUs was standardized and normalized to the minimum sequence number of samples (58). Alpha diversity indices such as Chao1 and Shannon indices were calculated for each sample using QIIME (version 1.7.0).

### The rust layer biofilm microbial and seawater microbial ecological process analysis

The microbial assembly of rust biofilm and seawater was influenced by the processes identified using mean nearest taxon distance (MNTD). To assess the level of non-random phylogenetic relatedness, we calculated the “normalized impact size” of the phylogenetic local area structure (ses. MNTD) by dividing the standard deviation of phylogenetic distances in the distribution by the difference between the phylogenetic distances obtained for the observed communities and null communities generated through 999 randomizations (59). These analyses were conducted using the “Picante” package in the R environment (Version 3.3.2). Specifically, the β-MNTD, which represents the mean distance between each taxon and its closest neighbor, was calculated by randomly rearranging OTUs and their abundances across phylogenetic tips to capture the uniqueness of bacterial networks (60). The term “β-NTI” refers to the difference between the mean of the null distribution and the observed “β-MNTD”. According to Vellend (61), the proportions of the influences of heterogeneous and homogeneous selection are determined by the scores of β-NTI values greater than or equal to 2. The Bray-Curtis distance (RCBray), was employed along with the β-NTI values to further quantify the contributions of major ecological processes shaping microbial assembly in rust layer biofilm and seawater. The fractions of homogenizing dispersal and dispersal limitation were assessed using β-NTI and RCBray values. Specifically, the portions of pairwise correlations with |β-NTI| < 2 and RCBray > 0.95 or < -0.95 were utilized to measure the overall significance of dispersal limitation or homogenizing dispersal processes individually (60). The score of pairwise comparisons with |β-NTI| ≥ 2 and |RCBray| ≥ 0.95 represented the contribution of ecological drift to compositional turnover (62). Selection and ecological drift exhibit deterministic or stochastic tendencies, respectively, among these processes, while dispersal can be either deterministic, stochastic, or both (63).

### Statistical analysis

The alpha diversities (Shannon index and Chao1 index) of bacterial and fungal communities in the rust layer biofilm and seawater were compared using Student’s t-test. To investigate the resemblance of microbial community structure between the rust layer biofilm and seawater, principal coordinate analysis (PCoA) was conducted utilizing the vegan package in R, based on Bray-Curtis distances (64). To determine if there were significant distinctions between the rust layer biofilm and seawater, an ANOSIM analysis was conducted using the R package (65). The ggplots package in R was utilized to generate a Venn diagram to identify shared and specific operational taxonomic units (OTUs) between the rust layer and seawater (66).

Through source tracking analysis and resolution, the contribution of marine bacterial community to the bacterial community in the rust layer biofilm was analyzed to clarify the influence mechanism of seawater bacterial community on the composition of bacterial community in the rust layer biofilm. Source Tracker analysis is used to identify different bacterial communities’ source of dissemination and estimate their contributions (such as relative abundance and dynamics of bacterial communities) (67). First, all samples are assigned with source or sink attributes; then, it is assumed that each sink sample is a mixture of source, the proportion of which is unknown; finally, the source composition proportion of each sample in the sink is obtained.

To evaluate species interplay of microbial communities in the rust layer and seawater, the top 50 most abundant genera were chosen for network construction via Spearman correlation coefficient ≥ |0.5| and p value < 0.05. A series of metrics were then computed, average path length, network diameter, average degree, clustering coefficient and graph density were calculated to analyze the topology of the network using Gephi platform (68).

## ACKNOWLEDGMENTS

This research was funded by the Guangxi Key Research and Development Program (Guike AB23026114) and the Guangxi Provincial Central Guidance Local Science and Technology Development Project (Guike ZY22096001-2).

Junxiang Lai, Congtao Sun, Maomi Zhao Jizhou Duan and Baorong Hou conceived and designed the research. Shengxun Yao and Zihan Pan performed the experiments and analyzed data. Shengxun Yao wrote the manuscript. All authors read and approved the manuscript.

The authors declare no conflict of interest.

## FUNDING

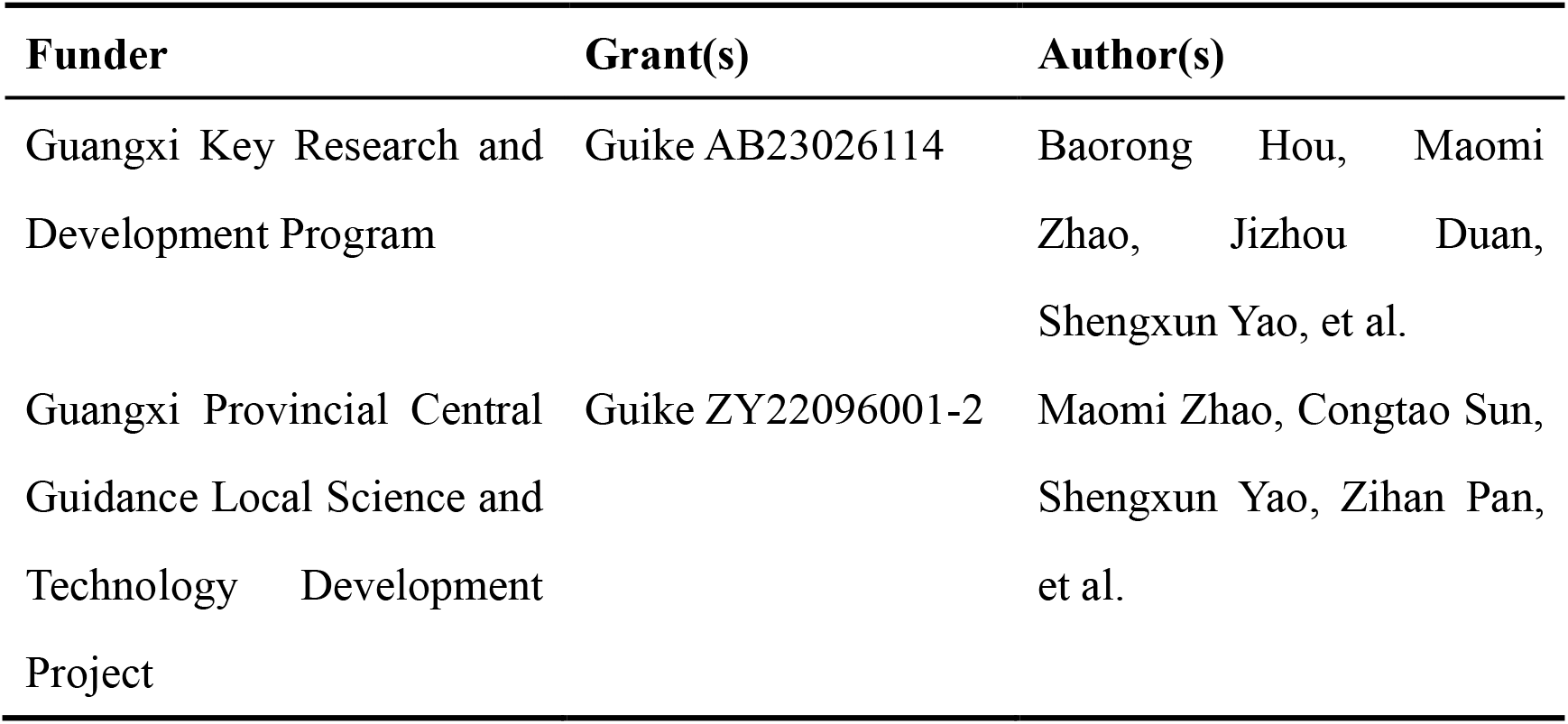

## DATA AVAILABILITY

All the sequences can be downloaded from the NCBI Sequence Read Archive Database under the accession numbers PRJNA1012345.

